# Characterizing enterotypes in human metagenomics: a viral perspective

**DOI:** 10.1101/2021.07.26.453761

**Authors:** Li Song, Lu Zhang, Xiaodong Fang

## Abstract

The diversity and high genomic mutation rates of viral species hinder our understanding of viruses and their contributions to human health. Here we investigated the human fecal virome using previously published sequencing data of 2,690 metagenomes from seven countries. We found that the virome was dominated by double-stranded DNA viruses, and young children and adults showed dramatic differences in their fecal enterovirus composition. Beta diversity showed there were significantly higher distances to centroids in individuals with severe phenotypes, such as cirrhosis. In contrast, there were no significant differences in lengths to centroids or viral components between patients with mild phenotypes, such as hypertension. Enterotypes showed the same specific viruses and enrichment direction after independent determination of enterotypes in various projects. Confounding factors, such as different sequencing platforms and library construction, did not result in a batch effect to confuse enterotype assignment. The gut virome composition pattern could be described by two viral enterotypes, which supported a discrete, rather than a gradient, distribution. Compared with enterotype 2, enterotype 1 had a higher viral count and Shannon index, but a lower beta diversity, indicating more resistance to the external environment’s harmful effects. Disease was usually accompanied by a viral enterotype disorder. However, a sample outside of the enterotyping mathematical space of enterotype database did not necessarily indicate sickness. Therefore, the background context must be carefully considered when using a viral enterotype as a biomarker for disease prediction. The disease, second only to the enterotype, explains significant variation in viral community composition, implying that double-stranded DNA is relevant to human health. Our results of investigating a baseline viral database highlight important insights into the virome composition of human ecosystems, and provide an alternate biomarker for early disease screening.

## 1 Background

In recent years, many studies have shown that viral colonization in the human body is highly related to human health and life. Cross-species virus transmission poses an extraordinary threat to human and animal health (Daszak et al., 2000). With advanced sequencing technology, the primary material for viral research has become viral genomes (virome), which enable viral identification and classification at the molecular level (Fujimoto et al., 2020; Gregory et al., 2020). The success of virome studies greatly relies on high-quality viral genomes (Minot et al., 2011). However, viruses are highly diverse and individual-specific (ref) and traditional purification strategies, culture, and sequencing are labor-intensive and inefficient (Reddy et al., 2015), thus severely preventing the comprehensive and intensive study of viruses.

The strategy of assembling the viral genome involves a comprehensive and in-depth analysis of the virome. David and colleagues launched the “Uncovering Earth’s virome” project to build The Integrated Microbial Genome/Virus (IMG/VR) database in 2016(Paez-Espino et al., 2016, 2017, 2019; Roux et al., 2021). Recently, data of 28,060 metagenomes were used to mine 142,809 human gut viruses, and Gubaphage was found to be the second common virus branch in the human gut(Camarillo-Guerrero et al., 2021). These projects opened the prelude to the construction of viral genome database and laid the foundation for a comprehensive analysis of the human gut virome(Gregory et al., 2020). Disease is associated with the gut virome, but studies have ignored the importance of viral sequencing information in massive metagenome sequencing data. The construction of the viral genome database has enabled detailed research on the human gut virome.

An enterotype is a cluster of microbes in the human gut and it describes the distributional of the human gut microbial community(Arumugam et al., 2011). Multiple studies have reported that there are two dominant enterotypes, which correspond to the individuals’ preference for digesting plant fiber or animal meat (Costea et al., 2017). The gut is an ecosystem, and the enterotype summarizes its microbial characteristics using mathematical methods(Arumugam et al., 2011; Holmes et al., 2012), but such knowledge is insufficient (Jeffery et al., 2012). Research on the composition patterns and function of the gut microbiome will significantly improve our understanding of its relationship with health and disease(Knights et al., 2014). Enterotypes can be used for gut microbial analysis, to inform disease treatment and prevention strategies, and may also provide a theoretical basis for diet therapy. The relationship between viral enterotypes and the human disease status is still largely unknown. Whether enterotypes can be used as biomarkers for predicting the disease status requires further research.

In this study, we collected previously published human metagenomic sequencing data, conducted sample quality control through a fast pipeline, identified virus species, and determined viral abundance. Furthermore, we established a baseline database of the human gut virome based on 2,690 metagenomes. We demonstrated the relationship between virus species and abundance in various ethnicities, countries, and diseases using different DNA library construction methods and sequencing platforms, and analyzed the association between viral community diversity and disease. Viral enterotypes were assigned by the Dirichlet multinomial mixture model (DMM). We independently identified enterotype-specific viral operational taxonomic units (vOTUs) for each dataset and resolved the inter-relationships among enterotypes from different projects by comparing the abundance of enterotype-specific viruses. Further, we compared the ecological diversity of viruses between different enterotypes, and evaluated the correlation of viral enterotype disorders and their diversity with diseases. The results of study elucidate the relationship between enteroviruses and human health in a large population and highlight the decisive role of viruses as molecular markers in identifying high-risk individuals. Viral research is likely to make an indispensable contribution to improving human health.

## 2 Materials and methods

### 2.1 Choosing an alignment method

Alignment and assembly methods are used to detect viruses and estimate their abundance. MetaPhlAn(Segata et al., 2012)and its upgraded version, MetaPhlAn2(Truong et al., 2015), are alignment tools that use marker genes for alignment and have achieved great success in bacterial genome alignment. However, many viral genomes do not have marker genes, and therefore, this strategy is not useful for viral classification and abundance estimation. Virome(Eric Wommack et al., 2012), VirSorter(Roux et al., 2015), and VirFinder(Ren et al., 2017) use assembly methods to classify viruses and calculate abundance, but these tools require a large amount of computing resources and time and therefore cannot be applied to large projects. Some recently developed alignment tools, such as ViromeScan(Rampelli et al., 2016), VIP(Li et al., 2016), and HoloVir(Laffy et al., 2016) have been shown to perform well for bacterial genomes. However, they are impractical for aligning viral reads to genomes. Moreover, many software tools are for online use, which means that they are unsuitable for large-scale projects. VirMap(Ajami et al., 2018) software developed for processing protein and genome data can provide good results. It can be accurately identify the virus species regardless of the sequencing depth. However, this software involves substantial computing resources. After comparing the advantages and disadvantages of different software(Ajami et al., 2018), we finally chose FastViromeExplorer(Tithi et al., 2018), a software based on k-mer alignment used by Kallisto(Bray et al., 2016). This software maps all reads to the reference and then uses the expectation-maximization algorithm to estimate the virus species and their corresponding abundance.

### 2.2 Data collection and processing

We downloaded all data from the National Center for Biotechnology Information (NCBI) sequence read archive (SRA). The SRA numbers for each project are shown in Supplementary Table 1. We only chose pair-end data from projects sequenced by the Illumina HiSeq 2000 or 2500 platforms. After processing the original data sample (Supplementary Figure 1) using Trimmomatics(Bolger et al., 2014)to remove the raw data and adapters of low-quality reads, we detected and removed contamination from the host’s DNA and RNA data, and discarded the unpaired reads. Finally, we used FastViromeExplorer software to align reads to IMG/VR v2.

### 2.3 Viral contig taxonomic annotation

We used Glimmer3 toolkit Version 3.02b(Delcher et al., 2007) to predict and extract the open reading frame of viral contigs with a minimum length threshold of 100 amino acids. The protein sequences were aligned to the UniProt TrEMBL database as of February 2021(Bateman et al., 2021) using BLASTX(Boratyn et al., 2012). The major voting system was then used as described previously to ascertain the family of a viral contig. A contig needed to be supported by five proteins to be considered as successful assignment; otherwise, the assignment was considered a failure. When a virus sequence was annotated to multiple families in taxonomic assignment, we choose the family with the largest proteins. When multiple families have the same number of proteins, the size of the accumulated E-value (BLASTX alignment) of all proteins was compared.

### 2.4 Calculation of ecological diversity

We first used Tximport(Soneson et al., 2015) R package to read the original abundance information of the virus (the output of Kallisto) from each project. The “betadiver” in the Vegan R package was used for calculating alpha and beta diversity. For alpha diversity, we first transformed the abundance information into integers and then used the “rrarefy” function to normalize abundance. We then used the “estimate,” “diversity,” and “specnumber” functions to obtain various measurement values of alpha diversity. We used the “RLE” method embedded in “calcNormFactors” to normalize raw abundance for beta diversity. We used the “Hellinger” method in “Decostand” to transform the data and eliminate false similarities caused by many viruses whose abundance was 0. When the abundance of many viruses in the two samples were 0, some algorithms might consider them to have similar abundance distribution and conclude that they were close to each other. The real reason may be that many viruses have not been detected. We used the “betadiver” and “betadisper” functions to obtain beta diversity, and then used Adonis2 to analyze the viral ecological differences between cases and controls, and corrected them with raw data size. The Kruskal test was used to determine whether there was a significant difference in the distance from the centroid between cases and controls. Tukey’s honestly significant difference test was used to determine differences in variance within and between groups.

### 2.5 Enterotyping and MaAsLin2 analysis

We used the DMM method to determine viral enterotypes in each project independently. Enterotypes were assigned using the “DirichletMultinomial “ R package, with predetermined parameters of 1 to 10 enterotypes, and enterotype data from each project were run 10 times. The smallest Laplace value corresponding to the number of enterotypes was considered as the optimal result. MaAsLin2 (huttenhower.sph.harvard.edu/maaslin2) analysis was used to determine the specific vOTUs associated with enterotypes, with correlations considered significant at the 5% level (after multiple testing correction). We also applied the envfit function in Vegan to estimate the effect size of the structural variance explained by factors such as enterotype and disease.

## 3 Results

### 3.1 Sequencing data and summarization

We collected 12.36 TB of metagenomic sequencing data from 18 previously published projects (Supplementary Tables 1 and 2). We selected data from 2,690 metagenome samples of high quality for the subsequent analysis (Supplementary Figure 1 and Supplementary Table 1), of which 1,092 were samples were from women, 859 were from men, and 739 were from unknown sex. The length of sequencing reads from each sample were 2.26 to 8.55 G (Supplementary Table 1), and approximately 10% of strictly filtered reads were aligned against IMG/VR v2 viral sequences (Supplementary Table 3). We obtained 2,690 metagenome samples by choosing paired-end sequencing data from the Illumina HiSeq 2000 and 2500 platforms and excluding projects with a small data size (< 1 G).

We annotated the geographic locations of the included projects on the basis of their predominant samples (Figure 1A). Because there were no specific sampling coordinates, each project was located by country. We annotated the viral taxonomy at the family level based on the protein sequence similarities (Minot et al., 2013; Hannigan et al., 2015). Approximately 50% of the viral genomes failed taxonomic assignment (Figure 1B), and double-stranded (ds) DNA viruses, such as Siphoviridae, Myoviridae, and Podoviridae, were the dominant enteroviruses as previously reported (Zuo et al., 2020). The density peak was close to zero, which indicated that the viruses were rarely shared among individuals (Supplementary Figure 2). The samples from Finland were outliers in the PCoA and tSNE plots (Figure 1C and D) because of the low viral diversity (Supplementary Figure 3). This finding might be explained by age. The average age of individuals in the Finland project was 1.5, and their gut communities did not reach stable states. Although the samples from the other six countries showed substantial variability in the PCoA and tSNE plots (Figure 1C and D), they belonged to the same cluster, especially the samples from the studies conducted in China. The studies from China had the most individuals, and the samples were spread over almost the entire plot. In the tSNE plot, we found that the samples from the USA and Peru were clustered in a local region, which indicated that the gut virome showed characteristics of geographical distribution.

**Figure 1:**
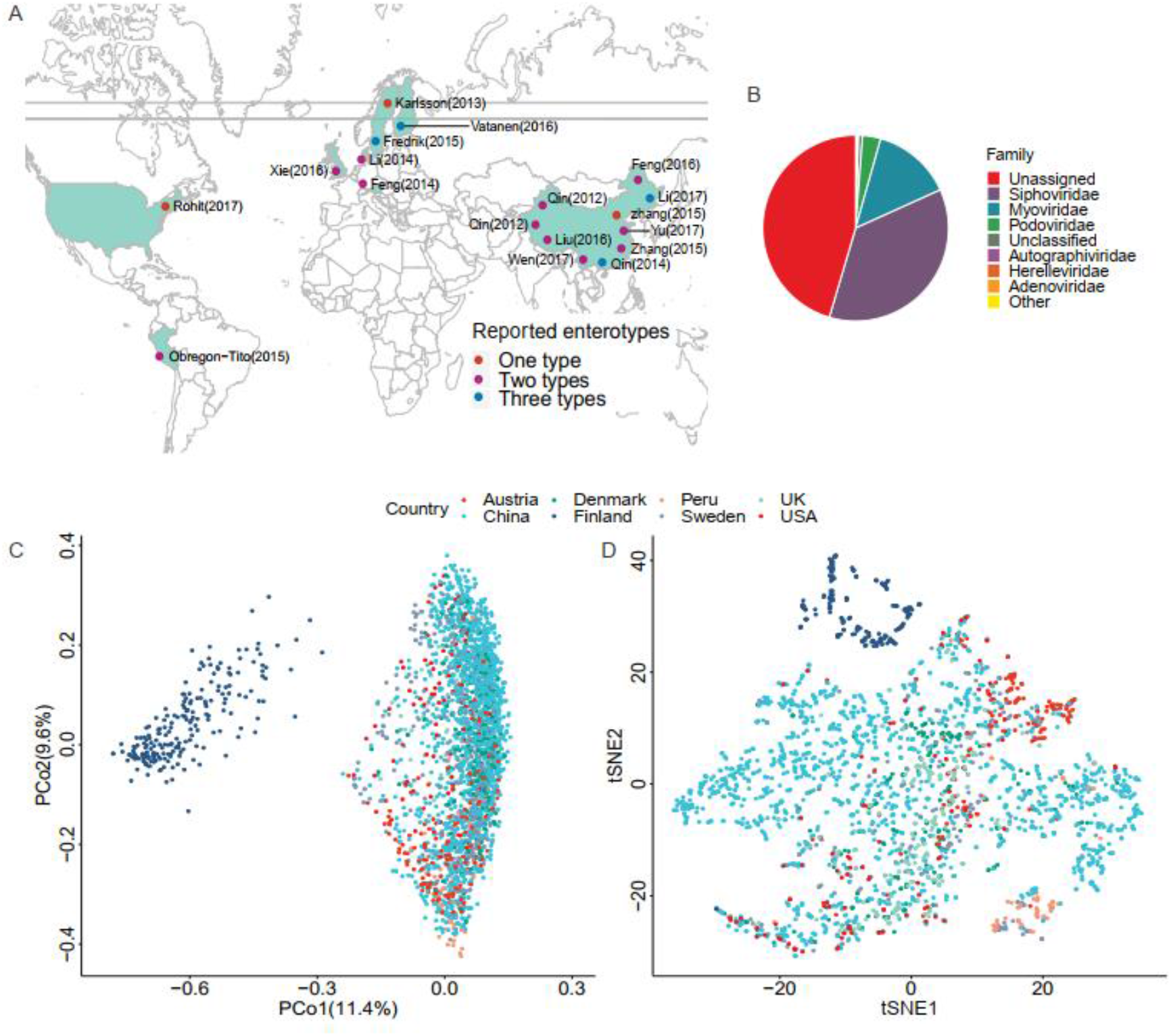
Location, taxonomic assignment, and abundance of the 2,690 samples. A: Geographic locations of the 18 projects, with classification by the number of enterotypes. B: Pie chart shows viral taxonomic assignment at the family level by protein alignment. C: Principal Coordinates Analysis (PCoA) plot based on the Bray–Curtis distance and the relative abundance of viruses. D: t - Distributed Stochastic Neighbor Embedding (tSNE) plot based on the relative abundance of viruses.

To study the distribution characteristics of the viral species in samples with different phenotypes, we divided all samples from studies with a case–control design into three categories. These categories of controls, cases, and all represented healthy people, patients with various diseases, and all individuals, respectively. As more samples were included, the number of viral species showed exponential growth, with no significant difference between cases and controls until samples from ∼100 individuals were included (Figure 2A). After including ∼100 individuals, the “case” curve showed a steep increased viral count. As expected, a significant increment in the number of viral species was observed when the number of samples was increased in the “all” curve. However, the three growth curves were essentially parallel (Figure 2A), which suggested that the overall number of viruses in the patient population after viral community disruption was limited. More interestingly, the “case” and “all” curves overlapped with each other after ∼1000 samples. The reason for this finding could be that the case population contained all species of viruses in the control population. When we compared the growth curves of different projects, we found that the curves for Finland, Peru, and Chinese populations with cirrhosis had significant differences (Figure 2B). The samples from the Finland project were obtained from only 1.5-year-old children, at which age the enterovirus community is not well established. It is unclear why the number of viral species in Peru samples were small at the beginning of the curve. The dramatic increase in the number of viruses in the Chinese population with cirrhosis may be due to severe disruption of the enterovirus community. We used unique species in cases and controls to define group-specific viruses and compared the change in the proportion of unique viral species between cases and controls (Supplementary Table 4). We found that the mean proportion of viruses in case samples was 26% and in control samples was 14%. Among all samples, the proportion of viruses that were unique to cases was 23%. Each case individual had an average of 10.99 viruses, and the ratio of viruses that were unique to controls was 4%, and each control individual had an average of 2.43 viruses. Overall, there was an enrichment of viruses in cases.

**Figure 2:**
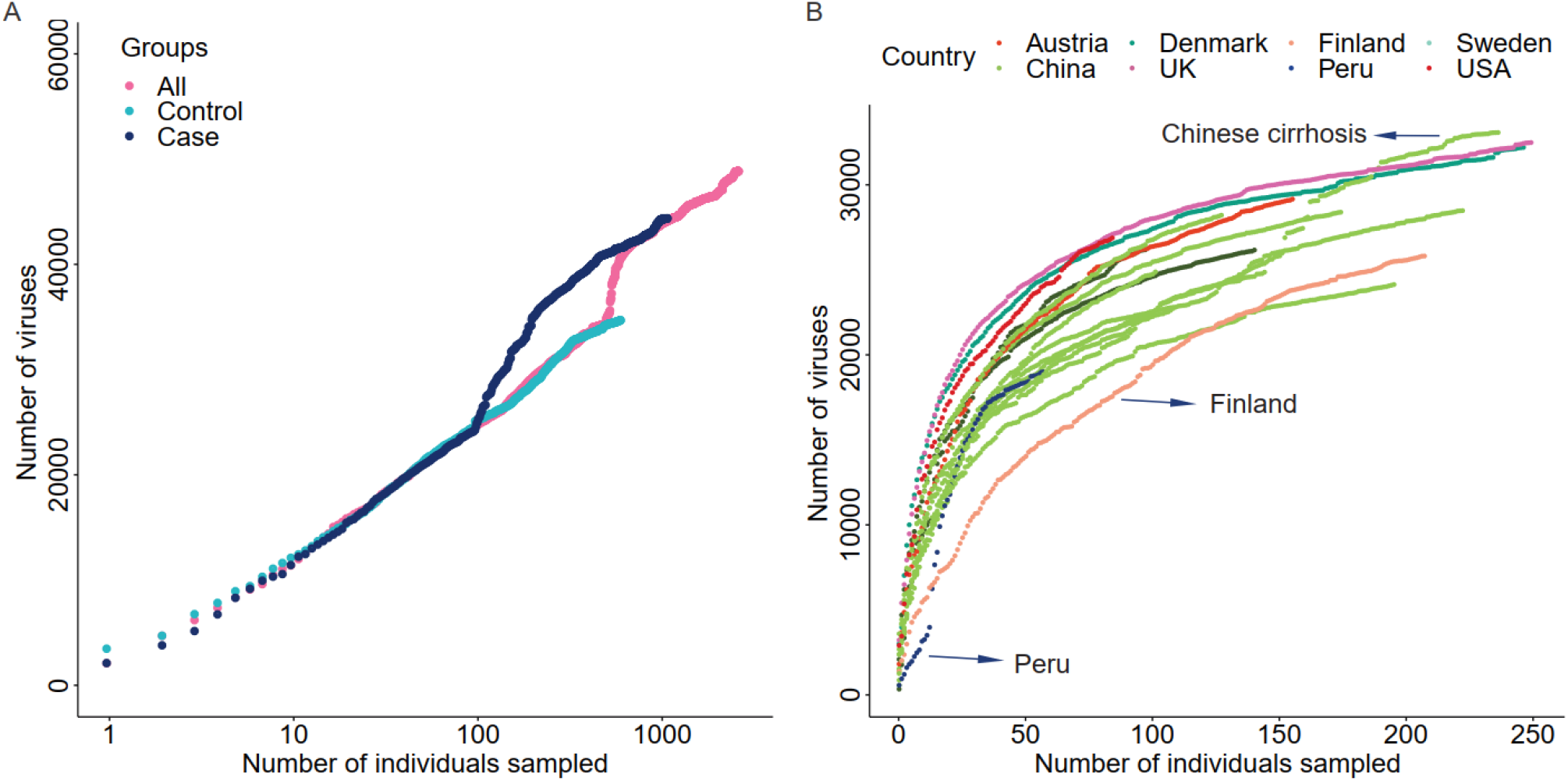
Cumulative curves of the number of virus species against the number of samples. A: Cumulative curves of cases, controls, and all samples. Only samples from studies with a case-control design were included. B: Cumulative curves of sample data divided into seven countries.

### 3.2 Relationship of ecological diversity of viruses and disease

The beta diversity of a microbial community is usually used to evaluate dynamic changes in an ecosystem (Koleff et al., 2003). A comparison of the results of projects with a case–control design revealed that the degree of imbalance in the viral community composition was related to the severity of the disease phenotype. An example of this finding is that the viral community in patients with cirrhosis (Figure 3A) was significantly different from that in healthy people (Adonis2, p = 0.001, adjusted for raw data size). Comparison of the distance to the centroid between patients and healthy individuals by the Mann–Whitney *U* test showed a significant dissimilarity (Figure 3B). Specifically, patients had a significantly larger distance than healthy individuals, which indicated that patients had a considerably disordered viral community. In contrast, we did not detect a significant difference between patients and healthy individuals in the hypertension project (Adonis2, p = 0.08, Figure 3C). We also compared the distance to the centroid for each pair of three cohorts (Figure 3D), and a significant difference was found only between patients with hypertension and healthy individuals (Wilcoxon, p = 0.036).

**Figure 3:**
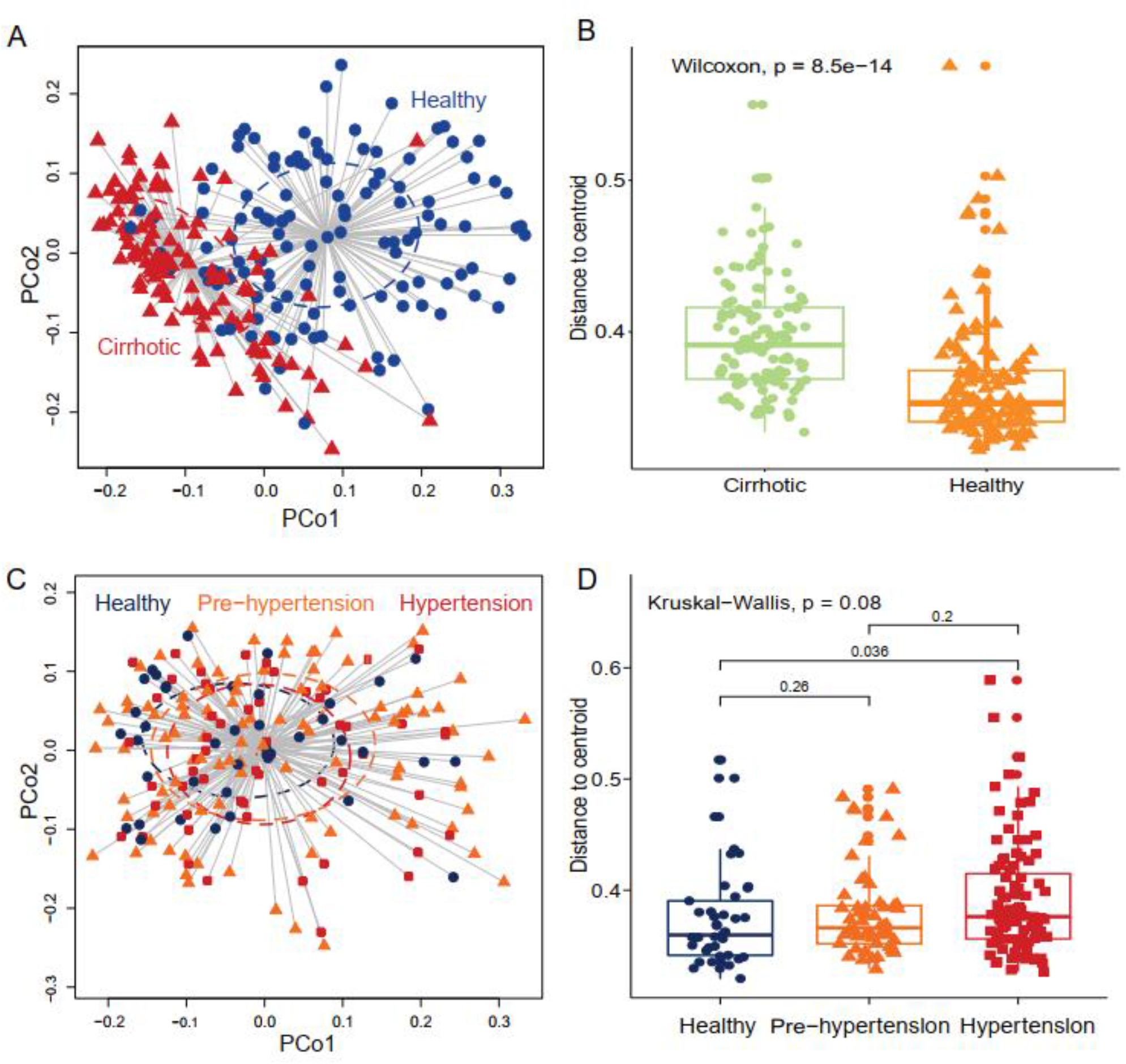
Gut virome characterized by beta diversity in the included projects. (A) Principal coordinates analysis plot of the cirrhosis project. Each ellipse represents a cohort, and the point connected by the straight gray lines represents the centroid. (B) Boxplot of the distance to the centroid. A significant difference in the distance to the centroid was found between the two groups. (C) Principal coordinates analysis plot of the hypertension project. (D) Boxplot of the hypertension project with comparison for each pair of the three groups.

We further investigated statistical differences in gut viral composition between case and control samples from various aspects to investigate changes in the viral community across different phenotypes. Using Adonis2, we found a significant difference in enteroviruses between cases and controls expect hypertension and obesity (Table 1), which suggested that their gut viral community was less affected by the disease state. Consistently, in cases with relatively mild phenotypes, such as hypertension or obesity, there was no noticeable differences in body metabolism compared with the controls. An analysis of variance (ANOVA) was used to determine whether there was a significant difference between two centroids (to test the component of viruses) between cases and controls. We found that the cirrhosis and cancer cohorts showed a substantial difference between two centroids (Table 1). The Kruskal-Wallis test was performed to determine whether the distance to the centroid in principal coordinates analysis was significantly different between the case and control groups, and the results were consistent with those of ANOVA. Compared with the controls, cases with more severe phenotypes, such as cirrhosis and cancer, showed substantial differences in gut viral composition (Table 1), whereas cases with relatively mild phenotypes, such as hypertension, showed no significant differences.

**Table 1.** Beta diversity for measuring the sample distance in projects with a case–control design.

| Project | Adonis2 for<br>disease | Adonis2 for<br>raw data | ANOVA | Kruskal<br>test |
| --- | --- | --- | --- | --- |
| Sweden T2D* | 5.00E-03 | 1.00E-03 | 6.34E-03 | 1.62E-03 |
| China cirrhosis | 1.00E-03 | 1.00E-03 | 3.91E-09 | 8.41E-14 |
| China rheumatoid arthritis | 2.40E-02 | 1.50E-02 | 0.86 | 0.48 |
| Austria carcinoma | 1.00E-03 | 0.55 | 5.29E-04 | 7.89E-05 |
| China colorectal cancer | 4.00E-03 | 3.10E-02 | 0.12 | 9.71E-03 |
| China hypertension | 0.08 | 0.25 | 0.13 | 0.08 |
| China coronary heart disease | 1.00E-03 | 0.81 | 4.15E-02 | 1.79E-02 |
| China T2D discovery | 2.60E-02 | 3.50E-02 | 1.75E-02 | 0.15 |
| China T2D validation | 1.00E-03 | 2.90E-02 | 0.38 | 0.45 |
| China obesity | 0.06 | 0.37 | 0.78 | 0.78 |

\*Type 2 diabetes (T2D).

### 3.3 Characterizing viral enterotypes

The characteristics of enterotypes of the gut virome were the focus of this study. Data on enterotypes are generally used to help adjust population stratification in Metagenome-wide association studies (MWAS) analysis (Wang et al., 2012). The correlation between enterotypes and disease phenotypes has received much attention in this field. The DMM method is commonly used for determining enterotypes of the gut microbiome and is more effective than the partitioning around medoids (Ding and Schloss, 2014). Different library construction methods, sequencing platforms, and other factors may lead to false-positive assignment of enterotypes. To avoid this situation, we adopted a project-independent strategy for determining enterotypes. There were two or three enterotypes in most projects, while some projects only had one enterotype (Figure 1A, Table 2, Supplementary Figure 4). Enterotypes with the same intrinsic composition pattern were considered as the same. We used Maaslin2 to discover enterotype-specific vOTUs and then determined their enrichment direction on the basis of mean abundance. Similar enterotypes had the same specific vOTUs and the same enrichment trend. We manually classified enterotypes in all of the projects into three groups (Table 2, Supplementary Table 5). Enterotypes 1 and 2, which are the two major types, were widely distributed in all projects, which indicated that these two types of enterotypes were common across the project populations. However, enterotype 3 was rare. Unclassified individuals were not able to be confidently assigned to enterotype 1 or 2.

**Table 2:** Manually categorized results for each project.

| Enterotype | Enterotype 1 | Enterotype 2 | Enterotype 3 |
| --- | --- | --- | --- |
| Denmark no phenotype | GP1 | GP2 | - |
| China cirrhotic | GP2 | GP1 | GP3 |
| Sweden mother-offspring pair | GP3 | GP2 | GP1 |
| China rheumatoid arthritis | GP1 | GP2 | - |
| Austria carcinoma | GP1 | - | GP2 |
| UK no phenotype | GP1 | GP2 | - |
| China colorectal cancer | GP1 | GP2 | - |
| China hypertension | GP2 | GP3 | - |
| China coronary heart disease | GP1 | GP2 | - |
| China T2D discovery | GP1 | GP2 | - |
| China T2D validation | GP1 | GP2 | - |
| China healthy Mongolian | GP1 | GP2 | - |
| China ankylosing spondylitis | GP1 | GP2 | - |

A permutation test was performed to demonstrate the validity of manual classification, which involved randomly paired enterotypes from different projects. We assumed that paired enterotypes had the same specific vOTUs and enrichment directions. We assigned a lower error rate to paired enterotypes if they had more identical vOTUs and similar enrichment trends. We repeated pairing 5 million times to obtain the distribution of pairing scores. These scores showed that our manually classified enterotypes had the lowest error rate (Figure 4A). Moreover, random pairing supported the three major enterotypes. Enterotypes 1- and 2-specific vOTUs were dominant (Figure 4B). The same enterotype-specific vOTUs with highly consistent enrichment trends indicated that the enterotypes from different projects had a similar pattern of virome composition (Figure 4B). Different DNA processing methods, sequencing platforms, ethics, age, and other confounding factors did not affect the identification of viral enterotypes. The vOTUs that were specific to unclassified enterotypes appeared complex. They intersected with either enterotype 1 or 2. Enterotype 3-specific vOTUs in different projects were less concordant than enterotypes 1- and 2-specific vOTUs.

**Figure 4:**
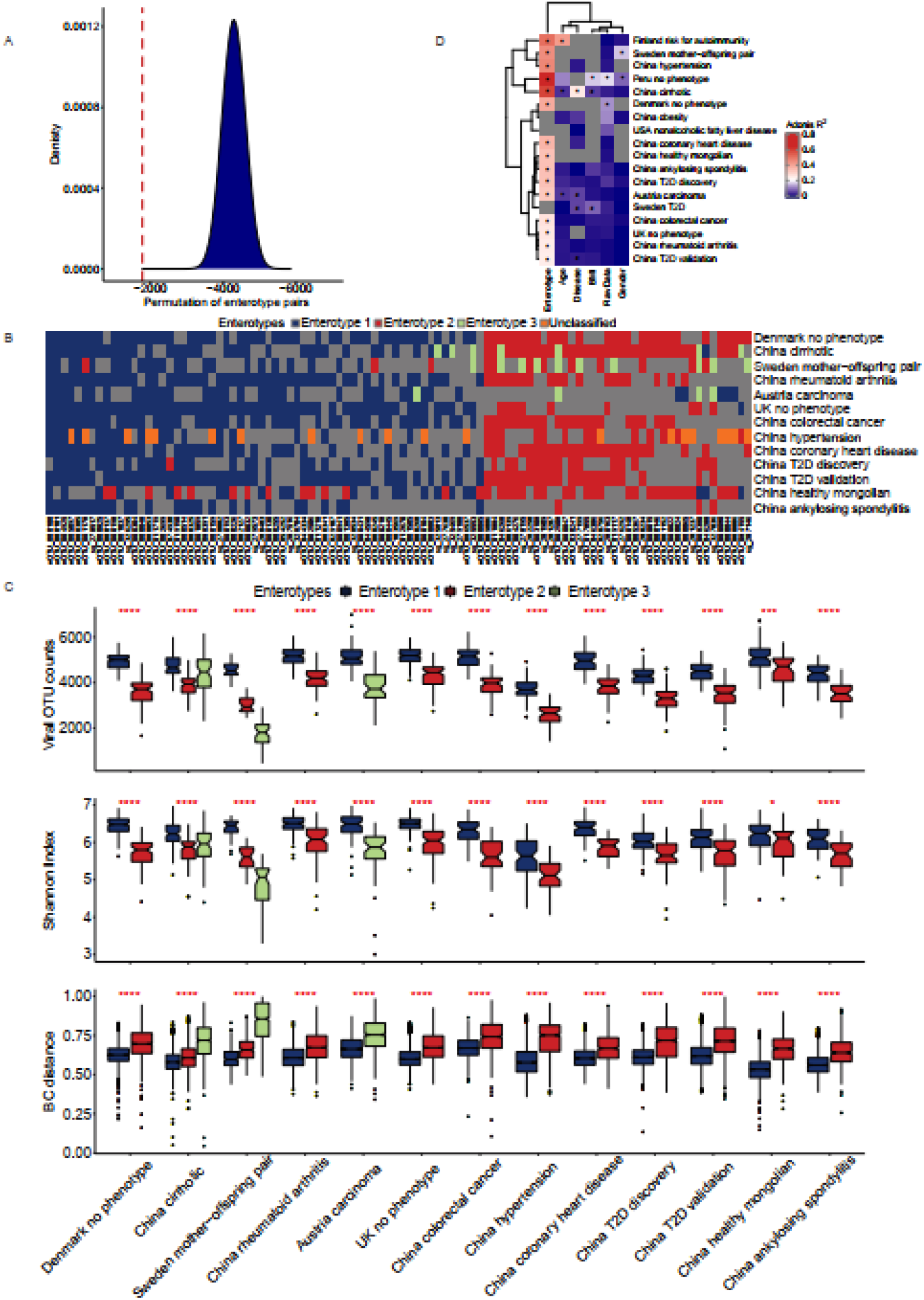
Characterization of viral enterotypes in all projects. A: We used the random pairing method to confirm the accuracy of artificial enterotype classification. The density map shows the score distribution of 5 million permutations, and the red line indicates the score of the manual category. B: The categories of manual enterotypes in different projects show a high concordance of their specific vOTUs and enrichment direction. C: Ecological diversity of different viral enterotype populations. D: Effect of different covariates on the structural variance of the gut virome community.

The microbiome is an ecosystem, the stability of which is reflected by the diversity of species in the system. As a species becomes more prosperous and uniform, the system’s diversity increases and it becomes more resistant to the effects of the external environment(Keesing et al., 2010). There are two dominant enterotypes in the viral community (Zuo et al., 2020), one of which has a high alpha diversity. The results of our study are remarkably close to expected results. Although the viral count varied among samples from different projects, enterotype 1 across the samples had more viruses than enterotype 2 (Figure 4C). A higher value of the Shannon index and a smaller sample distance in enterotype 1, compared with enterotype 2, indicated its more stable composition pattern. We found that more individuals were categorized as enterotype 1 than enterotype 2 (1204 vs. 716). By comparing the proportion of healthy samples with the two enterotypes, we found that individuals who were categorized as enterotype 2 had a higher risk of being sick than those who were categorized as enterotype 1 (odds ratio: 1.38, Fisher’s exact test, p = 0.01). We observed an interesting finding when we compared samples from the cirrhosis project and the Sweden mother-child project. The third enterotype had the most discrete sample distribution in the cirrhosis project, and a higher viral count and Shannon index compared with the Sweden mother-child project (Figure 3C). In contrast, the third enterotype had a large sample distance and the lowest viral count and Shannon Index in the Sweden mother-child project. The cases in these two projects had diverse medical conditions. Specifically, the case cohort in the cirrhosis project had disordered gut virome due to the disease, which explains why the number of viruses in the samples did not decrease. In contrast, children in the Swedish mother-child project lacked a stable gut virome and had a lower viral count, which suggested that enterotype 3 in the samples of this project was not caused by any disease.

The viral enterotype may play a dominant role in influencing the structural variance of the gut virome via a variety of factors. The Adonis test was used to determine the significance of viral enterotypes. The results were significant in all projects. Our results explain most of the structural variance in the gut virome (Figure 4D). In the Peru and cirrhosis projects, the Adonis R squared values were 0.62 and 0.57, respectively. Age, disease, BMI, raw data, and sex were not significant factors affecting viral enterotypes in most projects, but Adonis p values reached significance in several projects. Disease was the second most significant factor in the projects, which suggested that illness had a higher ability to reshape the gut microbiome than other factors. Characterizing the interaction between the gut virome and external stimuli was complex. Whether a single factor has a particular contribution requires consideration of the context of this factor. An example of this situation is that, in liver cirrhosis, the association between the gut virome and age was strong, but it was not significant for diabetes.

Groups in the same column were considered to belong to one enterotype.

Enterotypes are useful for describing the gut microbial community, and determining the association between diseases and enterotype is important to detect high risk individual in population. In the liver cirrhosis project, individuals could be broadly divided into three categories (Figure 5A). Enterotypes 1 and 3 were enriched in healthy individuals and patients, respectively (69 controls/16 cases vs. 2 controls/64 cases, Supplementary Table 6), and enterotype 2 accounted for half of them (43 controls, 43 cases, Supplementary Table 6). We found that the viral enterotype was significantly related to liver cirrhosis (Fisher’s exact test, p = 5.99E-24, Supplementary Table 7). Enterotype 3 was loosely distributed in individuals (Figure 5A). However, enterotypes 1 and 2 showed a closer relationship. These three groups did not have discrete clustering boundaries and demonstrated some overlap with one another in the PCoA plot. There was no apparent clustering of samples enriched locally due to the viral count or the Shannon index (Figure 5B). In the hypertension project, the clustering boundaries of enterotypes 1 and 3 were more pronounced than those for enterotype 2 (Figure 5C), and there was no overlapping area between the two clusters. This finding was surprising because individuals in enterotype 2 had a smaller viral count and a lower Shannon index (Figure 5D). Some of them were close to enterotype 1, while others had clusters of enterotype 3. However, the specific vOTUs and enrichment direction of individuals in enterotype 2 showed a high consistency (Figure 4B), indicating that enterotype 2 was real. We found no significant association between the viral enterotype and hypertension (Fisher’s exact test, p = 0.3, Supplementary Table 7). Gut virome community disorders showed significant differences in the cirrhosis and hypertension projects, which indicated that not all diseases caused evident ecological perturbation in the human gut. Thus, applying viral enterotypes as biomarkers for predicting clinical disease requires specific consideration.

**Figure 5:**
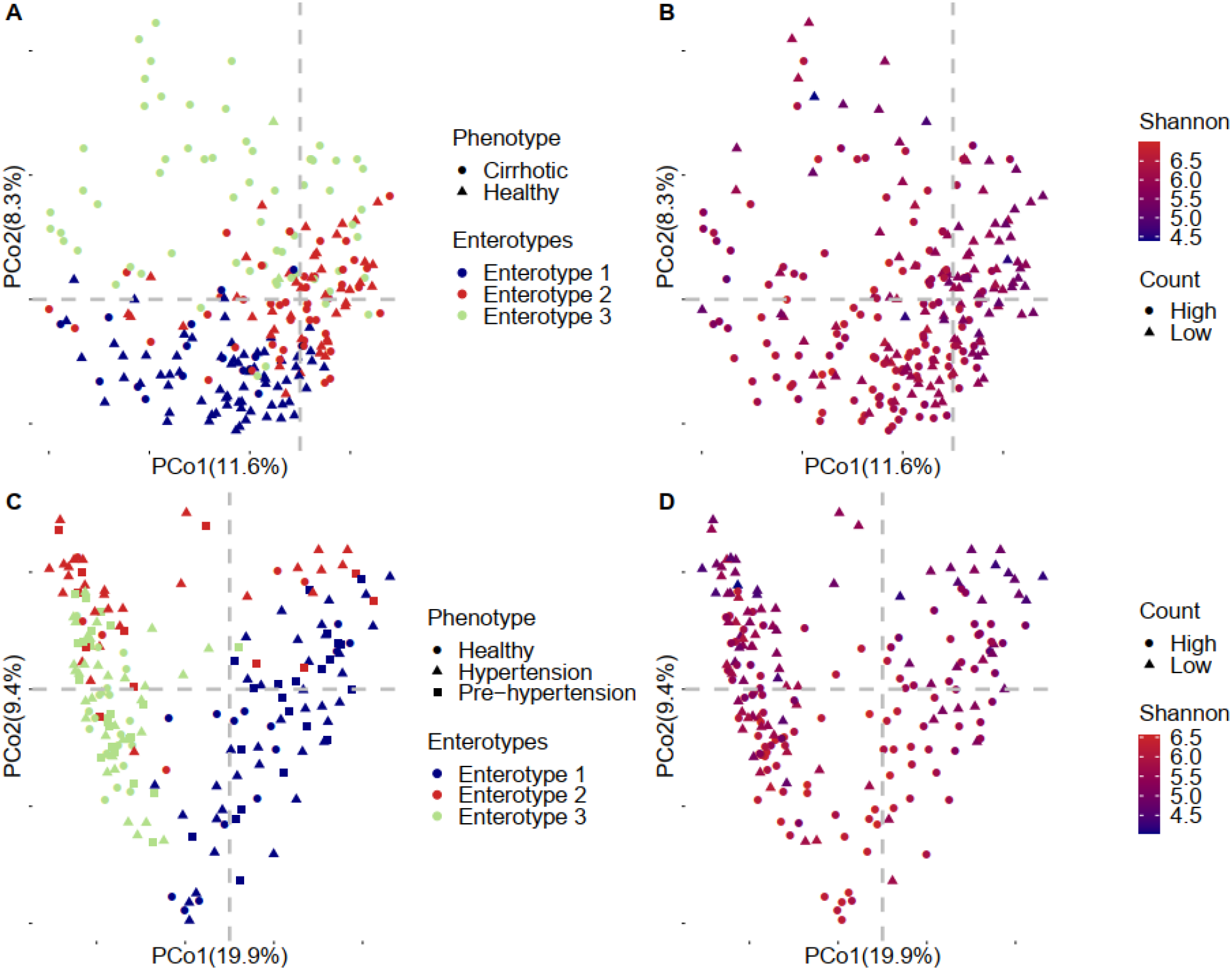
Detailed PCoA map of liver cirrhosis and hypertension. A: Samples of liver cirrhosis were plotted in relation to their phenotype and enterotypes. B: Samples of liver cirrhosis were plotted in relation to their viral count and Shannon index. C: Samples of hypertension were plotted in relation to their phenotype and enterotype. D: Samples of hypertension were plotted in relation to their viral count and Shannon index.

## 4 Discussion

Recent investigations have shown that enterotypes of the human gut can be divided into two categories based on their predominant flora (Bacteroidetes/Prevotella). Their functions correspond to the digestion of meat and vegetarian food (Arumugam et al., 2011; Costea et al., 2017). However, some researchers consider that the distribution of enterotypes is not discrete, but rather gradient. This viewpoint suggests that those two enterotypes are the two endpoints of the gradient distribution of Bacteroidetes/Prevotella(Jeffery et al., 2012). This study used the DMM method to assign viral enterotypes and showed that there were two enterotypes in most projects. Viral enterotypes did not have an apparent dominant virus. An explanation for this finding may be that most human enteroviruses are dsDNA viruses. As previously reported, dsDNA viruses are less harmful than RNA virus to the human body(Dutilh et al., 2014; Camarillo-Guerrero et al., 2021). These viruses do not undergo strong selection when colonizing the human gut, and there is no dominant viral strain that can occupy the whole human intestine. Recent studies have shown two common and harmless dsDNA virus branches in the human intestine, namely crAssphage(Dutilh et al., 2014) and Gubaphage(Camarillo-Guerrero et al., 2021). These two dominant virus branches may correlate with the two main viral enterotypes observed in this study. We analyzed the abundance of vOTUs corresponding to each enterotype and found that in different projects, the OTUs and enrichment direction of a specific virus in the same viral enterotype were consistent. Therefore, existing evidence and our findings support the view of two discrete viral enterotypes.

There was a third enterotype in several projects, but because of limited evidence, we could not conclude that it is ubiquitous in the human gut. In the hypertension project, researchers found that all specific vOTUs in the enterotype were shared with enterotypes 1 and 2. PCoA analysis also showed that individuals were located in the interconnection area, which is likely to explain a gradient distribution. In the liver cirrhosis project, 64 of the 66 samples were from patients, and the third enterotype was significantly related to patients. Unlike the hypertension project, the third enterotype in patients with liver cirrhosis was not related to the number of viruses. In other projects, viral enterotypes 1 and 2 had more specific viruses and a higher consistent enrichment direction in contrast to the rare specific viruses of viral enterotype 3 and an inconsistent enrichment direction. This result suggests that a third viral enterotype in various projects may not have belonged to the same cluster. Therefore, we cannot conclude that there was a stable presence of viral enterotype 3 in the population. We speculate that interaction between emergence of a disease and disorder of the gut virome may contribute to emergence of viral enterotype 3. Our results are in agreement with existing studies on the disruption of the human gut community that accompanies disease(Wang and Jia, 2016; Yu et al., 2017; Nakatsu et al., 2018).

In this study, enterotype 1 had a higher viral count and Shannon index compared with enterotype 2. In addition to having a smaller sample distance, we speculate that enterotype 1 might have more stable viral ecological communities than enterotype 2. We found enterotype 2 had 1.38 times more patients than the one in enterotype 1. This result suggests that a stable microbial community has a higher ability to resist the influence of external stimulations. In the cirrhosis project, enterotype 3 was characterized as being enriched in patients and having an extremely disordered gut virome. Enterotype 3 had the largest sample distance and highest number of virus species. A possible reason for this finding is that bile acid secretion in patients with liver cirrhosis is obstructed, which leads to drastic changes in the gut microbiome of the patients. This may have resulted in large-scale replacement of the virome and reduced similarity of virus species in this patient population. It is also possible that the microenvironment of viral evolution in the human body is disturbed owing to disease progression or the similarity of virus species is decreased due to a shift in the distribution of the ecological gradient. We also found a large distance in samples from the Sweden mother-child pair project, with the viral count being significantly lower than the global average level. A possible explanation for this finding is that the gut microbiome of young children is developing and has yet to reach a stable state(Derrien et al., 2019).

We determined enterotypes at the bacterial and viral levels in the China diabetes(Wang et al., 2012), and found a strong correlation between enterotypes at these two levels (China T2D discovery: p = 1.70E-07; China T2D validation: p = 1.58E-11, Fisher’s exact test, Supplementary Table 8). We found that enterotypes (bacterial and viral levels) were not randomly distributed and that the bacterial community had a strong selection effect on the viral community. However, bacterial- and viral-level enterotypes were not correlated or were weakly correlated with sex, age, BMI, and disease (Supplementary Table 9). This finding may be explained by the use of high-abundance bacterial and viral species for determining enterotypes. A high abundance of bacteria or viruses in the intestine is significantly related to disease. Such a high abundance directly and severely affects the human body, That is a microbial infection, not a harmonious symbiosis, which is in contrast to the current understanding of the gut community and health. Wang et al. used enterotype as a covariate in their study and proposed a useful method to stratify human gut microbiomes in MWAS, which effectively improved the power of hypothesis testing(Wang et al., 2012). Although we found a strong correlation between bacterial and viral enterotypes, they were not proven to be equivalent. Therefore, we suggest using bacterial and viral enterotypes as independent covariates in MWAS. More in-depth investigations are warranted to determine whether this strategy can efficiently reduce false-positive and false-negative rates of investigating pathogenic microbiomes.

We used MaAsLin2 to identify viruses that were specific to enterotypes and disease. We found that the number of vOTUs that were specific to disease was significantly lower when we simultaneously entered these two factors into the software program than when we entered only disease. In the cirrhosis project, 56 vOTUs were specific to the disease state (q-value ≤ 0.05), when the enterotype was excluded. In contrast, we found 241 and 7 vOTUs were specific to the enterotype and disease, respectively, when we included these two factors simultaneously. There were 21 vOTUs tested as disease-related became enterotype-associated. These results suggest that viral enterotypes need to be taken into account is MWAS. Although much of the literature suggests that most dsDNA viruses are not strongly associated with disease, we cannot rule out the contribution of dsDNA viruses to illness.

To better determine the efficacy of using enterotypes to identify the disease state, a standardized method is first required for determining enterotypes. One such standardized pipeline for enterotypes was previously reported by Costea et al. (2017). Researchers have also established an enterotype database of the gut microbiome community on the basis of the MetaHIT dataset (Qin et al., 2010; le Chatelier et al., 2013). They then built a machine learning model and trained it by applying the MetaHIT dataset and the corresponding enterotypes. Finally, this model was used to predict the enterotypes of testing samples on the basis of their bacterial abundance matrix. We independently assigned enterotypes in different projects and found that the two manually adjusted categories shared the most specific viruses and similar enrichment directions. This consistency masked the batch effects among different datasets, and demonstrates the ability of viral enterotypes to identify individuals with disease. The construction of a large-scale viral enterotype database to define the enterotyping mathematical space of healthy individuals might be helpful to detect individuals with disease outside the mathematical space. Therefore, we believe that using viral enterotypes of the gut virome community as a feature for disease prediction will significantly improve the accuracy of disease prediction.

## 5 Author Contributions

XF conceived this study. LS, LZ, and XF analyzed data, prepared the figures, and drafted the manuscript.

## 6 Funding

This work was financially supported by the Science Technology and Innovation Committee of Shenzhen Municipality, China (SGDX20190919142801722).

## 7 Conflict of Interest Statement

The authors declare that the research was conducted in the absence of any commercial or financial relationships that could be construed as a potential conflict of interest.

## 8 Acknowledgments

The authors thank many interns and former colleagues for collecting data, and their colleague Yufen Huang for discussing the analysis strategy.

## Notes

### Competing Interest Statement

The authors have declared no competing interest.

