## Supplementary figures and images for "Characterizing enterotypes in human metagenomics: a viral perspective"

### Supplementary Figure 1

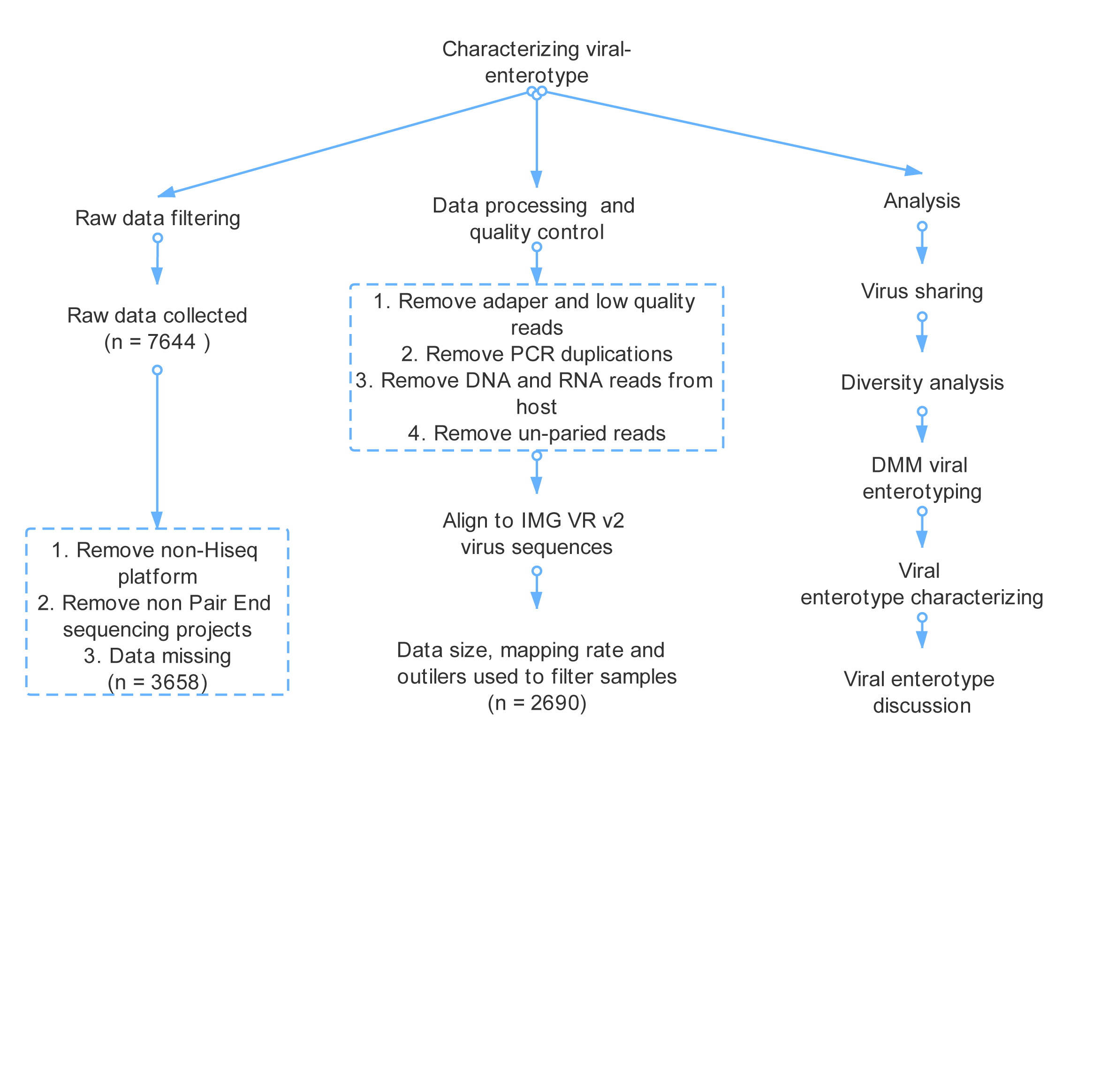

### Supplementary Figure 2

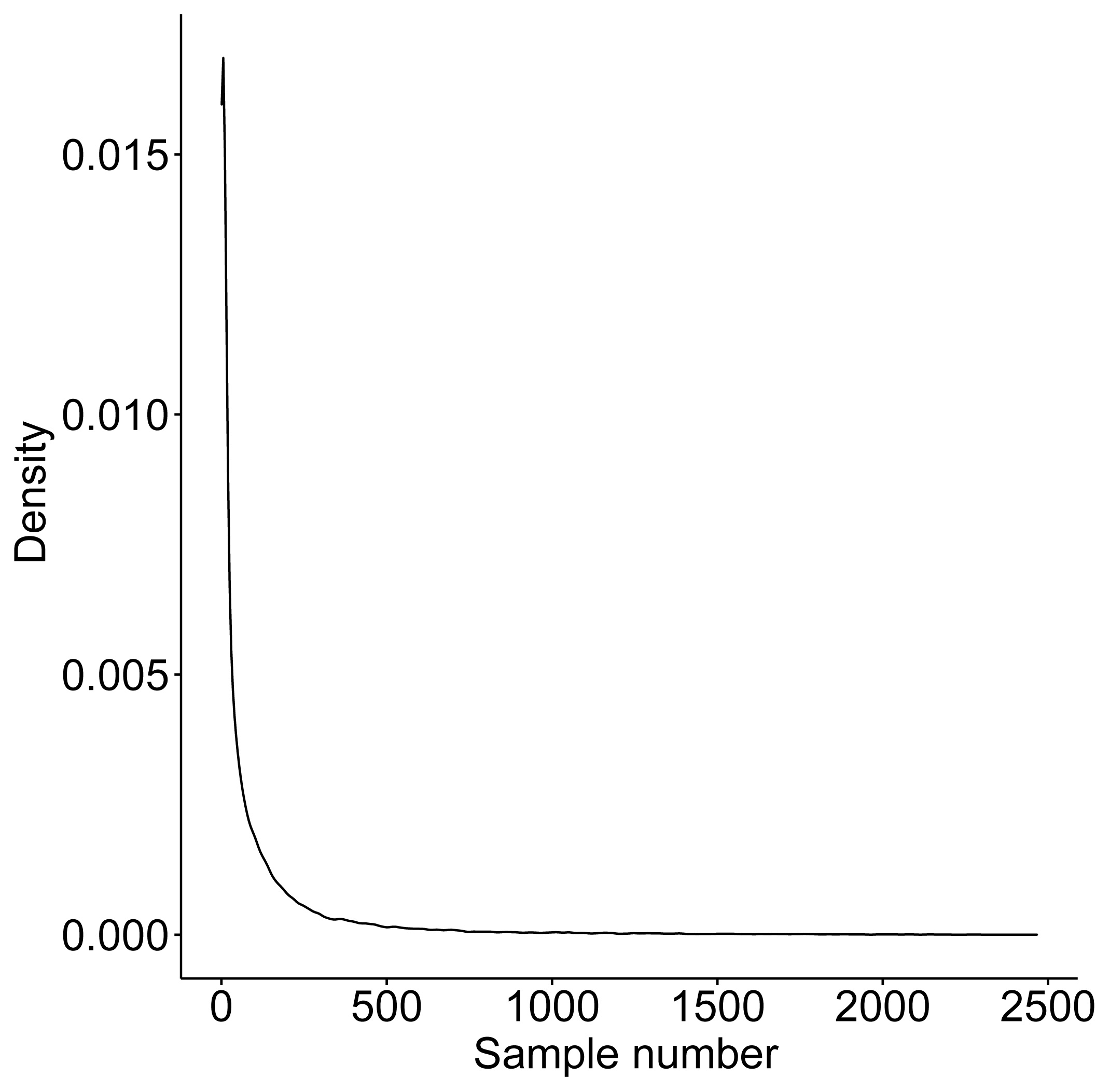

### Supplementary Figure 3

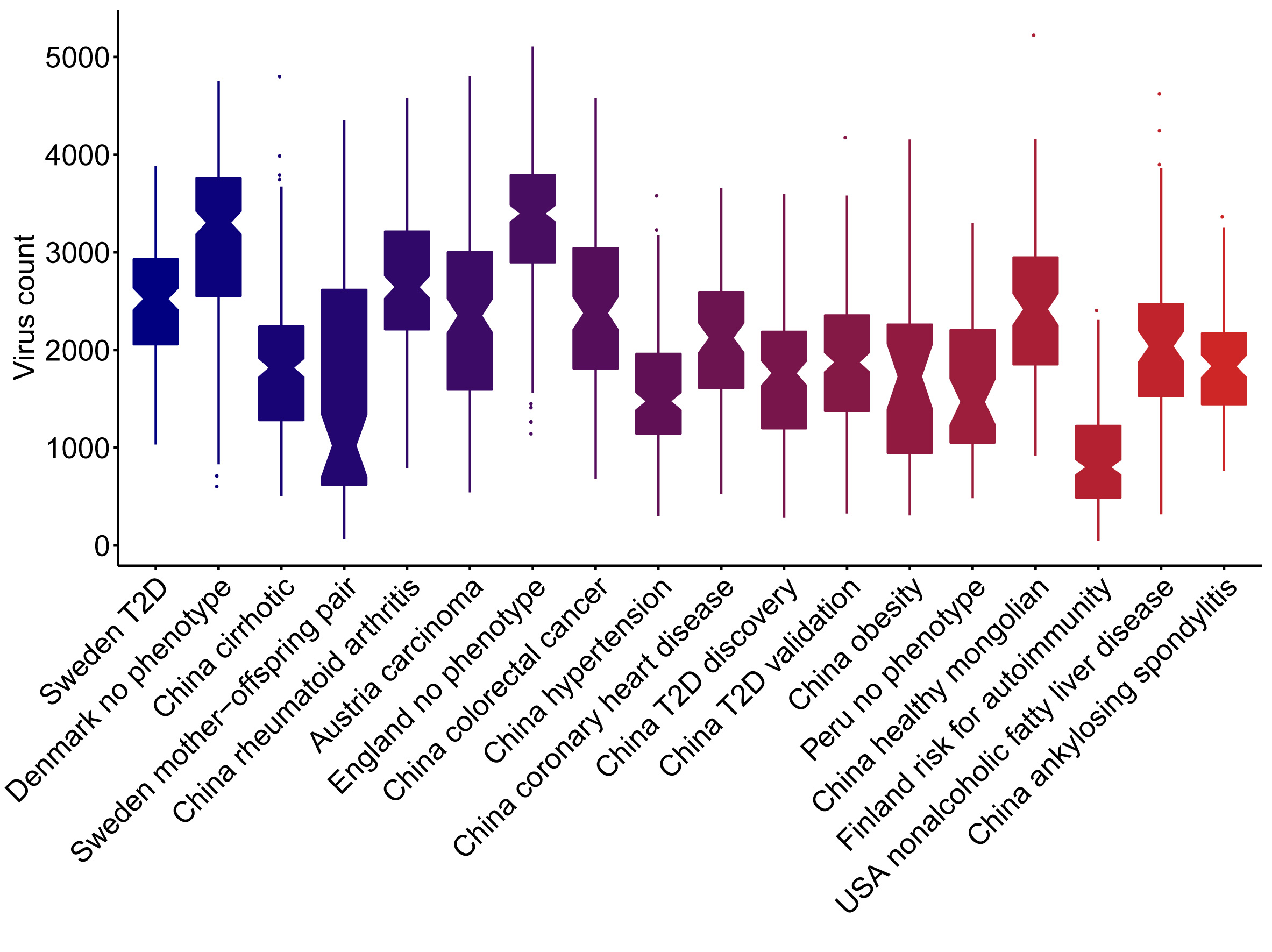

### Supplementary Figure 4

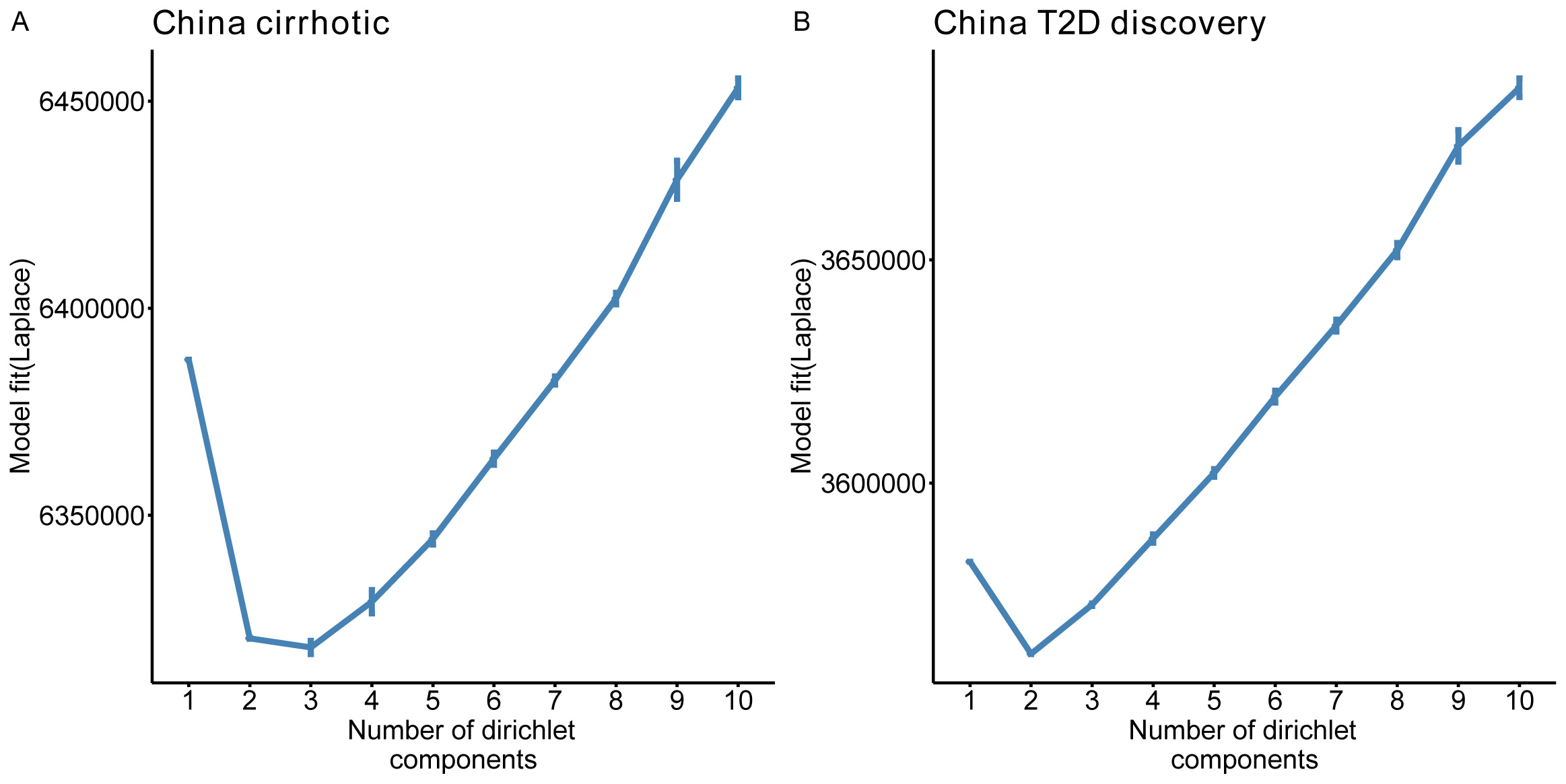
